# Understanding monocyte-driven neuroinflammation in Alzheimer’s disease using human brain organoid microphysiological systems

**DOI:** 10.1101/2025.02.16.638539

**Authors:** Chunhui Tian, Zheng Ao, Jonas Cerneckis, Hongwei Cai, Lei Chen, Henyao Niu, Kazuo Takayama, Jungsu Kim, Yanhong Shi, Mingxia Gu, Takahisa Kanekiyo, Feng Guo

## Abstract

Increasing evidence suggests that Alzheimer’s disease (AD) pathogenesis strongly correlates with neuroinflammation. Peripheral monocytes are crucial components of the human immune system that may play a role in neuroinflammation, but their contribution to AD pathogenesis is largely understudied partially due to the lack of appropriate human models. Here, we present human cortical organoid microphysiological systems (hCO-MPSs) for modeling dynamic AD neuroinflammation mediated by monocytes. By incorporating 3D printed devices into an existing cortical organoid protocol, 96 hCO-MPSs can be established with significantly reduced necrosis and hypoxia as well as enhanced viability within a commonly used 96 well plate, and each hCO-MPS consists of a doughnut-shaped hCO and a 3D printed device per well. Using this approach, monocytes from AD patients exhibit higher infiltration, decreased amyloid-beta (Aβ) clearance, and stronger inflammatory responses compared to monocytes from age-matched control donors. Moreover, pro-inflammatory effects such as elevated astrocyte activation and neuronal apoptosis were observed to be induced by AD monocytes. Furthermore, the significant increase in the expression of IL1B and CCL3, both at the transcriptional and protein levels, indicated the pivotal role of these cytokine and chemokine in monocyte-mediated AD neuroinflammation. Our findings provide insight for understanding monocytes’ role in AD pathogenesis, and the user-friendly MPS models we present are compatible with existing laboratory settings, highlighting their potential for modeling neuroinflammation and developing new therapeutics for various neuroinflammatory diseases.

## Introduction

Alzheimer’s disease (AD) is the most common form of dementia that affects an estimated 44 million people worldwide, and is characterized by a progressive loss of memory and cognitive function^1^. Increasing clinical evidence suggests that neuroinflammation plays a critical role in AD pathogenesis^2–4^, as shown by neuroinflammatory changes in AD patients through molecular imaging^5, 6^ and postmortem brain staining^7, 8^. While the excessive inflammatory activation of brain-resident immune cells in response to AD-related neuropathology is well-documented and accelerates neurodegeneration, the role of the peripheral immune system in AD-related neuroinflammation and neurodegeneration is just beginning to be understood. Peripheral monocytes, a crucial component of the peripheral immune system, are increasingly recognized for their importance in inflammation^9, 10^, sparking significant interest in their role in AD neuroinflammation and potential impact on disease progression. Endpoint analyses of postmortem AD brain tissue show extensive monocyte infiltration into the brain parenchyma of AD patients, where they cluster around amyloid-beta (Aβ) plaques^11, 12^. The population of infiltrated monocytes were also found to be increased as the progression of AD worsens^11^.However, dynamic interactions of monocytes and brain parenchyma remain poorly understood in AD neuroinflammation, partly due to the absence of suitable human models.

Human brain organoids (hBOs) are three-dimensional (3D) *in vitro* brain-like tissues derived from human stem cells and hold a remarkable potential for modeling and understanding neurological disorders^13–16^. With their 3D structures and cell components resembling the human brain, hBOs can replicate the complex brain microenvironment^13–15, 17^, a capability not achievable by 2D in vitro models. Meanwhile, being entirely derived from human cells, hBOs can fully replicate human biology and potentially reduce the translational barriers from experimental findings to therapeutic applications^18–20^, offering advantages over animal models^21, 22^. In recent years, the hBO technology has been extensively applied to model AD-related pathology and uncover how complex cell-cell interactions deteriorate as AD progresses^23^. Brain organoids derived from genetically edited human pluripotent stem cells (hPSCs) or AD patients induced hPSCs (hiPSCs) have been generated to study risk gene factors such as APP, PSEN1/2, and APOE4 mutations^24–27^. Additionally, human brain organoids derived from healthy hiPSCs and human embryonic stem cells (hESCs), have been co-cultured with microglia^28, 29^ or exposed to various non-cellular factors, such as Aβ^30^ and AD serum^31^, to model various AD-related pathologies. However, current human brain organoids still hold their power in studying AD neuroinflammation due to their limitations including high heterogeneity, low throughput, necrosis and hypoxia, lack of immune components, etc. Recently, microfluidics and organ chip advances have been employed to engineer better organoids using sophisticated microfabricated device and complicated engineering systems^32–34^. There is a tremendous need to develop simple, robust, scalable, and user-friendly organoid models and engineering platforms for wide applications in common research lab settings.

Herein, we developed innovative human cortical organoid microphysiological systems (hCO-MPSs) for understanding monocyte-driven AD neuroinflammation. By incorporating 3D printed devices into an adapted cortical organoid protocol, we can engineer and culture 96 hCO-MPSs within a commonly used 96 well plate, and each MPS consists of a doughnut-shaped hCO and a 3D printed device. Our MPS models may possess several unique aspects. Firstly, inspired by the neural tube structure, we engineer MPSs with the doughnut-shaped hCOs to model AD neuroinflammation. This unique doughnut shape design can not only enhance oxygen/nutrient diffusion to eliminate the necrotic and hypoxic conditions, but also facilitate the incorporation of immunocytes (e.g., monocytes), compared to conventional spheroidal hCOs. Secondly, through incorporating with time-lapse imaging, our MPSs can enable the tracking of dynamic immune-organoid interaction in 3D human brain-like microenvironments in parallel. Additionally, compared to current microfluidic and organ chip approaches, our MPSs are simple, robust, and scalable, due to the employment of simple 3D printing technology. Finally, our MPSs are user-friendly and compatible with current organoid protocols that use well plates and orbital shakers in common research lab settings. Thus, our MPS models allow us to study dynamic immune-neuron interaction in a 3D human brain-like environment, especially to understand monocyte-mediated AD neuroinflammation.

## Results

### Human cortical organoid microphysiological systems (hCO-MPSs) for the study of monocyte-mediated neuroinflammation in AD

To model interactions between monocytes and brain components during AD neuroinflammation, we developed human cortical organoid microphysiological systems (hCO-MPSs) which integrate AD monocytes and doughnut-shaped hCOs (**Fig. 1a**). AD monocytes, derived from the peripheral blood mononuclear cells (PBMCs) of AD patients, carry patient-specific disease information. When integrated with cortical organoids generated from human iPSCs which provide a 3D human brain microenvironment, the MPS allows us to investigate monocyte-mediated AD neuroinflammation.

**Figure 1.**
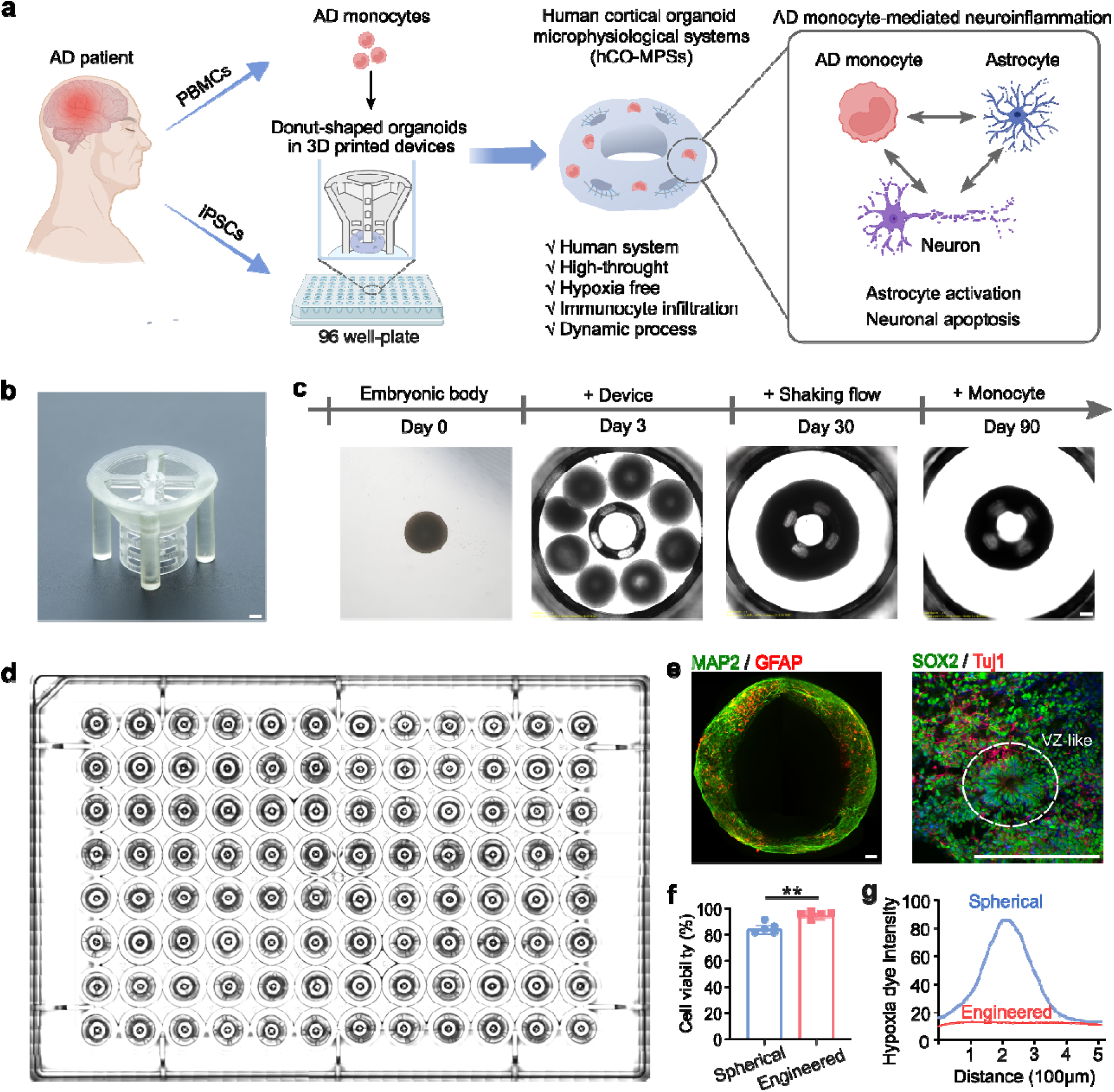
Human brain organoid microphysiological systems (hCO-MPSs) fo understanding monocyte-mediated neuroinflammation in AD. **(a)** Schematics of modeling monocyte-mediated neuroinflammation in AD using hCO-MPSs. **(b)** The image of a 3D printed device for generating doughnut-shaped hCOs. Scale bar: 500 μm (**c**) A timeline showing doughnut-shaped hCO generation in the MPS system. Scale bar: 200 μm. (**d**) The 96-well-compatible hCO-MPS platform for doughnut-shaped cortical organoid differentiation. **(e)** Whole mount staining of 3-month-old doughnut-shaped hCOs for GFAP (a mature astrocyte marker) and MAP2 (a mature neuron marker) (left) and 1-month-old doughnut-shaped hCOs for Tuj1 (an early-stage neuron marker) and SOX2 (a neural progenitor marker) (right). The white dashed line indicates the ventricular zone (VZ)-like structure. Scale bar: 100 μm. (**f)** Quantification of cell viability in three-month-old spherical and engineered doughnut-shaped hCOs cultured on the MPS device (meanL±Ls.e.m, n = 5 organoids from three independent experiments). (**g)** Quantification of hypoxia dye intensity in three-month-old spherical and engineered doughnut-shaped hCOs.

Inspired by the drought-like cross section of neural tubes, which are necrosis and hypoxia free, we used unified 3D-printed scaffolds compatible with the 96-well culture format to derive doughnut-shaped hCOs in a high-throughput manner (**Fig. 1b**). The device for generating doughnut-shaped hCOs is comprised of two main components: (1) a hollow circular scaffold with a mesh structure that allows for efficient culture medium perfusion and monocyte loading into the hCO (**Fig. S1**); and (2) a round glass bottom that holds pluripotent stem cell-derived embryoid bodies (EBs), facilitating the formation of hCOs. The EBs grow in the space between the perfusable scaffold and the bottom glass, gradually fusing to form doughnut-shaped hCOs (**Fig. 1c & S2**). Doughnut-shaped hCOs contain mature microtubule-associated protein 2 positive (MAP2^+^) neurons and glial fibrillary acidic protein positive (GFAP^+^) astrocytes (Fig. 1d, left) as well as exhibit ventricular zone (VZ)-like tissue organization (**Fig. 1d, right**). Furthermore, the doughnut-liked shape as well as the shaking flow and the perfusable scaffold promote efficient nutrient and oxygen exchange, leading to significantly increased hCO cell viability on-device culture (**Fig. 1e & S3a**). Specifically, doughnut-shaped hCOs have an average ring width of 0.5 mm (**Fig. S3b & S3c**), effectively preventing hypoxia and necrosis in organoids (**Fig. 1f**). In contrast, a significant portion of spherical hCOs that have an average radius of 0.9 mm (**Fig. S3b & S3c**) cannot be efficiently supplied with oxygen and nutrients due to diffusion constraints, resulting in the formation of an obvious hypoxic core (**Fig. 1f**). Overall, we engineered a microphysiological system that supports efficient hCO differentiation and maturation, resolves hypoxia, and can be easily scaled for high-throughput experiments.

### Infiltration study of monocytes from AD patients and age match controls

We then aimed to create a physiologically relevant platform of monocyte infiltration in AD by using monocytes isolated from PBMCs of AD patients and age-matched control (AC) donors **(Table S2)**. Before introducing AD monocytes into the system, we first incorporated the THP-1 monocyte cell line to assess the feasibility of our hCO-MPSs (**Fig. S4**). The results highlighted the superior 3D brain microenvironment provided by the organoids (**Fig. S4a & 4b**), and the successful infiltration of THP-1 cells, especially in the condition that simulated the activated state of AD monocytes and the AD brain environment (LPS/INF-γ pre-stimulated THP-1 cells in Aβ-pretreated doughnut-shaped hCOs) (**Fig. S4c-e)**. AD monocytes were then introduced into the hCO-MPSs to mimic monocyte infiltration into AD patient brains. Time-lapse images captured over 24 hours revealed gradual AD monocytes (red) infiltration into the hCO-MPSs (green, **Fig. 2a & Supplementary Movie 1**). Compared to conventional spherical hCOs, nearly twice as many monocytes infiltrated hCO-MPSs, highlighting the advantage of using our hCO-MPS platform to model monocyte infiltration **(Fig. 2b & S3e)**. Enhanced monocyte infiltration into hCO-MPSs can be attributed to the higher surface area-to-volume ratio of doughnut-shaped hCOs compared to spherical hCOs, thus increasing the contact area for monocytes **(Fig. S3d).**

**Figure 2.**
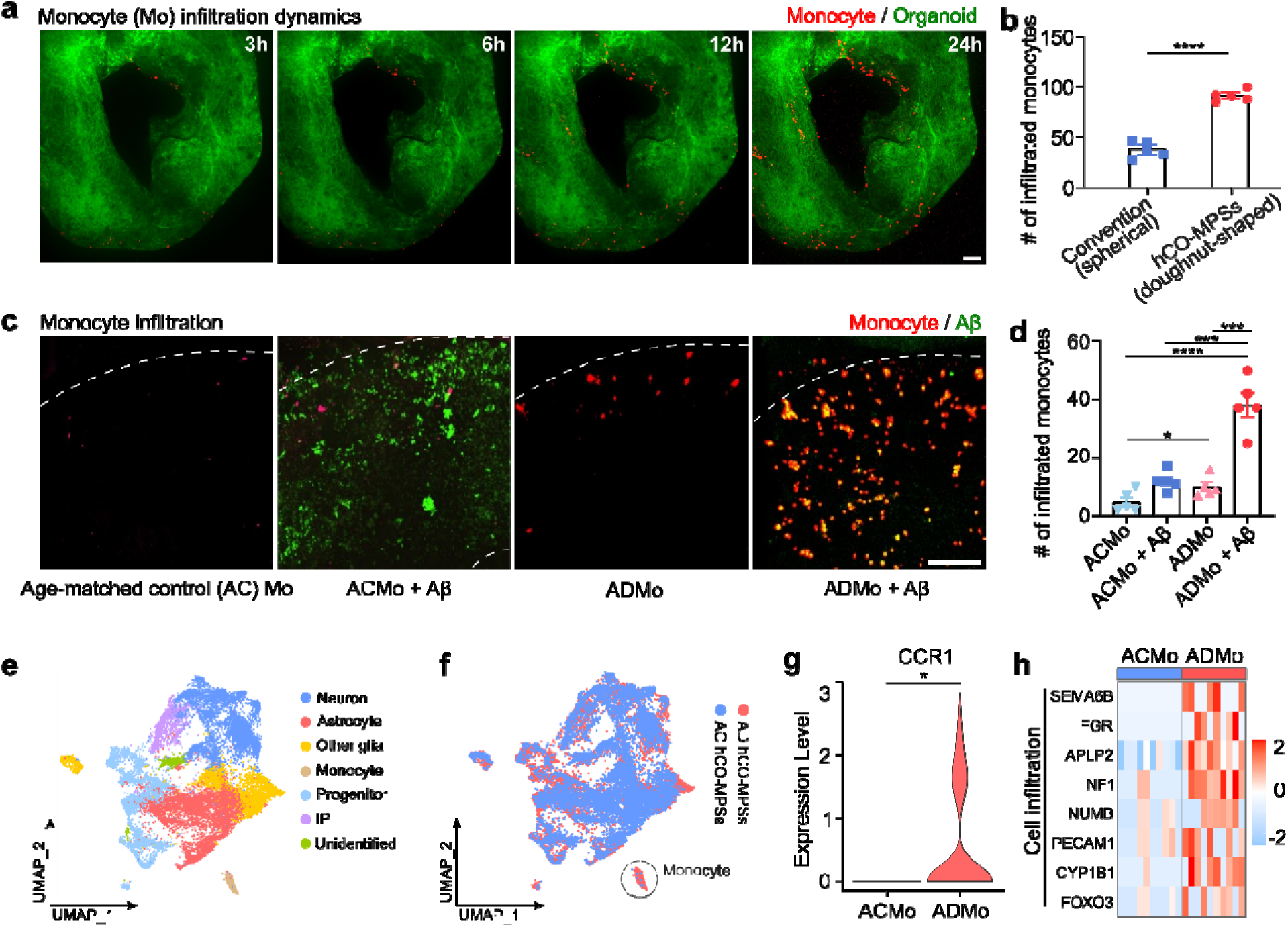
Infiltration study of monocytes from AD patients and age match controls. **(a)** Dynamic infiltration of AD monocytes (ADMo) into the hCO-MPSs over 24 hours. **(b)** Quantification of AD monocyte infiltration into conventional spherical hCOs and the hCO-MPSs with doughnut-shaped hCOs over 24 hours (mean ± s.e.m., n = 5 organoids from three independent experiments). **(c)** Representative images showing monocyte infiltration into the hCO-MPSs under different conditions: age-matched control monocytes (ACMo), ACMo + Aβ, ADMo, and ADMo + Aβ. **(d)** Quantification of monocyte infiltration experiment shown in (c) (mean ± s.e.m., n = 5 organoids from three independent experiments). **(e)** UMAP visualization of cell types in ACMo and ADMo hCO-MPSs. **(f)** UMAP visualization of the cell distribution of ACMo and ADMo hCO-MPSs. **(g)** Gene expression of CCR1 in ACMo and ADMo within hCO-MPSs. **(h)** Expression of cell infiltration-related genes in ACMo and ADMo within hCO-MPSs. **(a, c)** Scale bar: 100 μm.

We next asked if monocyte infiltration into hCO-MPSs is influenced by the disease status and AD-related neuropathology. To this end, we quantified AC monocyte and AD monocyte infiltration into hCO-MPSs. Interestingly, we found that AD monocyte infiltration into hCO-MPSs was significantly higher than that of AC monocyte, suggesting that monocytes in the AD patient blood possess cell-intrinsic properties that promote their infiltration into the brain tissues **(Fig. 2c & 2d)**. Furthermore, Aβ treatment that mimicked AD brain environment substantially increased AD monocyte but not AC monocyte infiltration into hCO-MPSs, underscoring the role of Aβ in peripheral monocyte infiltration in AD patients. AD monocytes also co-localized with Aβ in the hCO-MPSs (**Fig. 2c**), supporting the observations that infiltrated monocytes accumulate around Aβ plaques in the brain tissues of AD patients^11^. To study the mechanisms of increased AD monocyte infiltration into the hCO-MPSs, we performed single-cell RNA sequencing (scRNA-seq) of AD monocyte-enriched hCO-MPSs (AD hCO-MPSs) and AC monocyte-enriched hCO-MPSs (AC hCO-MPSs) **(Fig. 2e & Fig. S5a)**. Cell clustering confirmed higher infiltration of AD monocytes compared to AC monocytes **(Fig. 2f)**. We detected increased expression of CCR1, an important receptor for monocyte recruitment, as well as a number of other cell migration-related genes including SEMA6B, FGR, APLP2, NF1, NUMB, PECAM1, CYP1B1, FOXO3^35–40^ in AD monocyte as compared to AC monocytes **(Fig. 2g & 2h)**. In summary, the functional *in vitro* experiments and gene expression changes both revealed the distinct increased infiltration of AD monocytes into brain tissue, which further enhanced in the presence of Aβ.

### Altered functionality of AD monocytes in hCO-MPSs

Given our observation that AD monocytes co-localize with Aβ debris, we next investigated whether these monocytes can effectively phagocytose Aβ and thus assist brain-resident cells in clearing AD-related debris^41, 42^. Upon closer inspection of monocyte-infiltrated hCO-MPSs, we detected an extensive overlap of Aβ (green) and monocyte (red) signals in AC monocytes, but the overlap was decreased in AD monocytes **(Fig. 3a & 3b)**. These findings indicate that although monocytes can readily clear Aβ deposits, their clearing capacity is decreased in AD. Indeed, we found that the expression of phagocytosis-related genes, including *FCER1G*, *FCGR2A*, *TYROBP*, and *GAS6*^43–46^, was downregulated in AD monocytes as compared to AC monocytes **(Fig. 3c)**, indicating a reduced ability of AD monocytes to phagocytose Aβ. Similarly, several genes reported to be involved in Aβ catabolism, such as *RAB5A*, *RAB11A*, *CLU*, and *LRP2*^47–51^, were also downregulated in AD monocytes, suggesting the impaired Aβ catabolism in AD monocytes. Meanwhile, among the differentially expressed genes (DEGs) between AD and AC monocytes in hCO-MPSs, we found that *S100A8* and *LYZ* were the most upregulated genes in AD monocytes **(Fig. 3d)**. Given their major roles in the immune response and inflammation^52, 53^, the significant upregulation of *S100A8* and *LYZ* suggested an enhanced inflammatory response in AD monocytes. Consistent with our *in vitro* findings, Gene Ontology (GO) enrichment analysis confirmed pathways related to elevated inflammatory response and cell migration, as well as reduced phagocytosis and Aβ metabolism in AD monocytes (**Fig. 3e & S5b**). Overall, we found the altered functionality of AD monocytes in hCO-MPSs, including decreased impaired Aβ clearance ability and increased inflammatory response.

**Figure 3.**
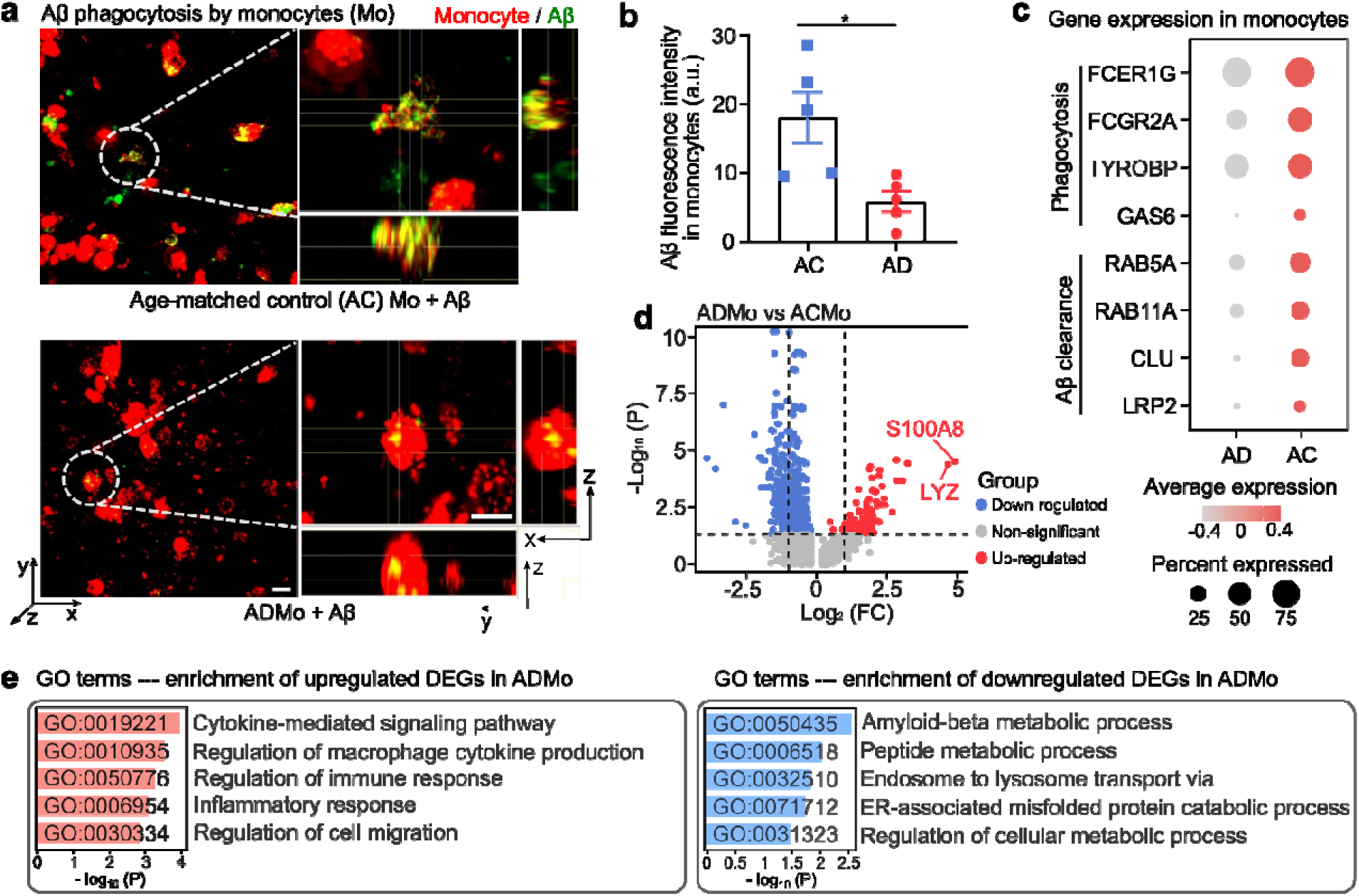
Functional characterization of AD monocytes (ADMo) and age-matched control monocytes (ACMo) in hCO-MPSs. **(a)** Representative images showing Aβ phagocytosis by ACMo and ADMo within hCO-MPSs. Scale bar: 10 μm. **(b)** Quantification of Aβ phagocytosis shown in (a) (mean ± s.e.m., n = 5 organoids from three independent experiments). **(c)** Expression of phagocytosis- and Aβ catabolism-related genes in ACMo and ADMo within hCO-MPSs. **(d)** A volcano plot displaying differentially expressed genes (DEGs) in ACMo and ADMo within hCO-MPSs. **(e)** Gene Ontology (GO) enrichment analysis based on ADMo DEGs.

### AD monocytes induce astrocyte activation in hCO-MPSs

Having observed enrichment for inflammatory pathway activation in AD monocytes, we hypothesized that infiltrating peripheral monocytes induce astrocyte activation in the brain tissue of AD patients, which is a well-described pro-inflammatory phenomenon that contributes to AD progression^54, 55^. To evaluate astrocyte activation in AD and AC monocytes, we first performed immunostaining for GFAP since reactive astrocytes are known to exhibit elevated *GFAP* expression and enlarged cell bodies **(Fig. 4a & S6)**^56^. We also quantified the production of reactive oxygen species (ROS), as astrocytes are a major source of ROS in the brain and their production increases upon astrocyte activation^57, 58^. AD hCO-MPSs exhibited significantly higher GFAP fluorescence intensity and enlarged area. As well as increased ROS production, as compared to AC hCO-MPSs, which were further increased in the presence of Aβ **(Fig. 4b-d)**. These findings support our hypothesis of heightened neuroinflammation under AD pathology. In agreement with our functional experiments, astrocyte gene expression analysis revealed upregulation of inflammatory and ROS-related genes. GO enrichment analysis also indicated an elevated inflammatory response in AD hCO-MPSs as compared to AC hCO-MPSs **(Fig. 4e, 4f & S5c)**. Meanwhile, enriched processes among the downregulated DEGs suggested impaired astrocyte ability to support neurons in AD hCO-MPSs **(Fig. 4f)**.

**Figure 4.**
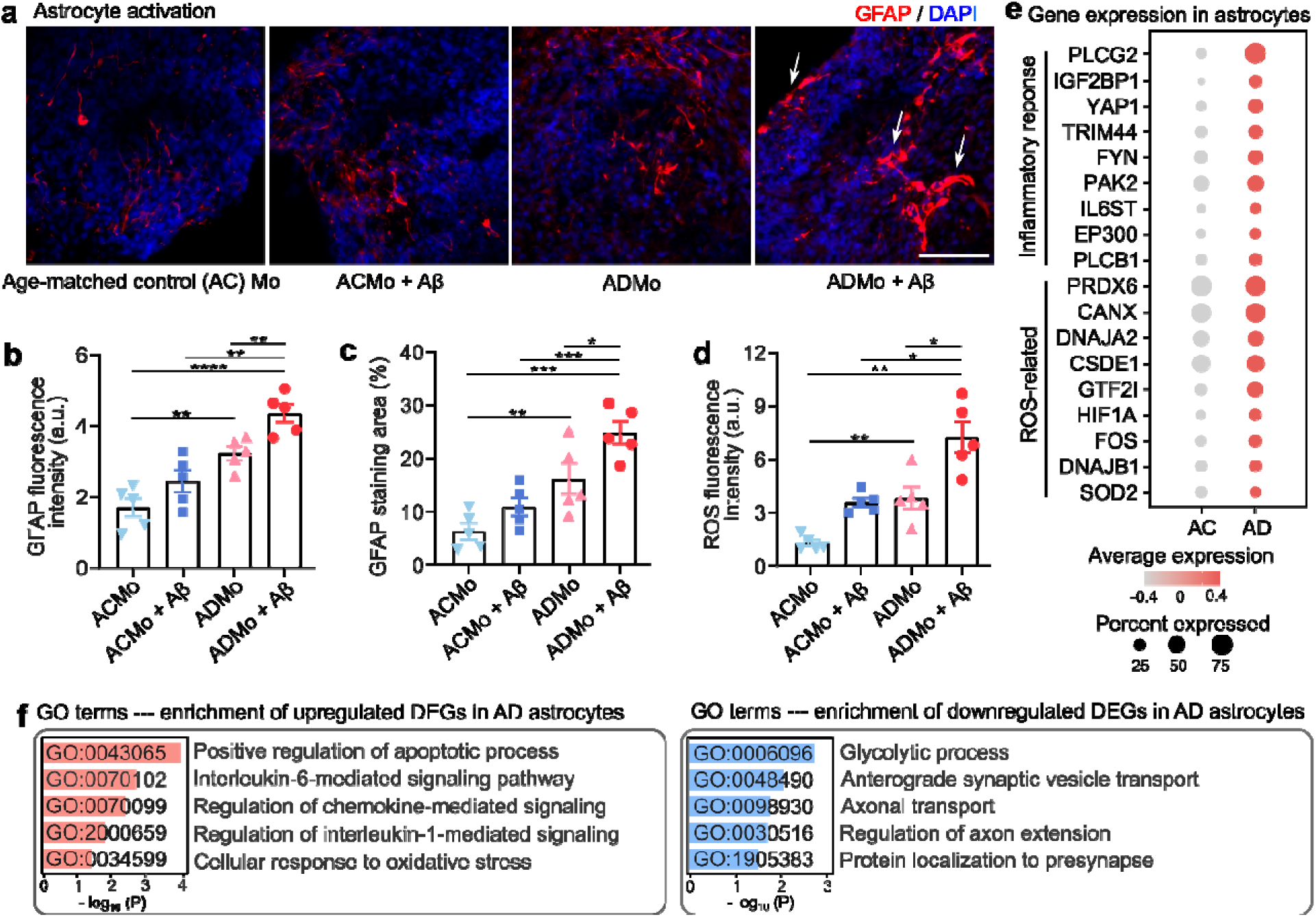
AD Monocytes (ADMo) induce astrocyte activation in hCO-MPSs. **(a)** Representative images showing astrocyte activation in hCO-MPSs under different conditions: age-matched control monocytes (ACMo), CMo + Aβ, ADMo, and ADMo + Aβ. Scale bar: 50 μm. **(b)** Quantification of GFAP fluorescence intensity for (a) (mean ± s.e.m., n = 5 organoids from three independent experiments). **(c)** Quantification of GFAP^+^ area for (a) (mean ± s.e.m., n = 5 organoids from three independent experiments). **(d)** ROS levels in hCO-MPSs under different conditions: ACMo, ACMo + Aβ, ADMo, and ADMo + Aβ (mean ± s.e.m., n = 5 organoids from three independent experiments). **(e)** Expression of immune response- and ROS-related genes in astrocytes within ACMo and ADMo hCO-MPSs. **(f)** Gene Ontology (GO) enrichment analysis based on astrocyte DEGs in ADMo hCO-MPSs.

### AD Monocytes induce neuronal apoptosis in hCO-MPSs

Cellular AD pathologies eventually converge on neuronal network degeneration and neuron cell death that led to cognitive decline^59, 60^. Neuroinflammation may cause neuron cell death by disrupting the supportive microenvironment of the human brain^61, 62^. Therefore, we investigated whether infiltrating monocytes could induce neuronal apoptosis in hCO-MPSs. Within 24 hours of AD monocytes co-culture with hCO-MPSs, we observed a gradual destruction of the ventricular zone (VZ)-like structure accompanied by cell vacuolation, indicating the detrimental effects of AD monocytes on the brain tissue **(Fig. 5a & Supplementary Movie 2)**. We performed the terminal deoxynucleotidyl transferase dUTP nick end labeling (TUNEL) assay to quantify neuronal apoptosis in hCO-MPSs **(Fig. 5b & S7**, apoptotic neurons are indicated by white arrows). TUNEL analysis revealed significantly higher neuronal apoptosis in AD hCO-MPSs as compared to AC hCO-MPSs, which was further increased in the presence of Aβ **(Fig. 5c)**. The expression of cell apoptosis-related genes *ATF4*, *JUN*, *GADD45G*, *ITGB1*, *RPS3*, and *ANP32A*^63^ was also significantly increased in neurons within AD hCO-MPSs as compared to AC hCO-MPSs, supporting the *in vitro* observations **(Fig. 5d)**. Finally, GO analysis revealed enrichment for apoptosis, DNA damage, and stress response processes among upregulated DEGs as well as synapse organization, neuron development, and neuron projection extension processes among downregulated DEGs **(Fig. 5e & S5d)**. Taken together, these findings indicate that AD monocytes can induce neuronal apoptosis and thus may contribute to neuronal network degeneration in AD.

**Figure 5.**
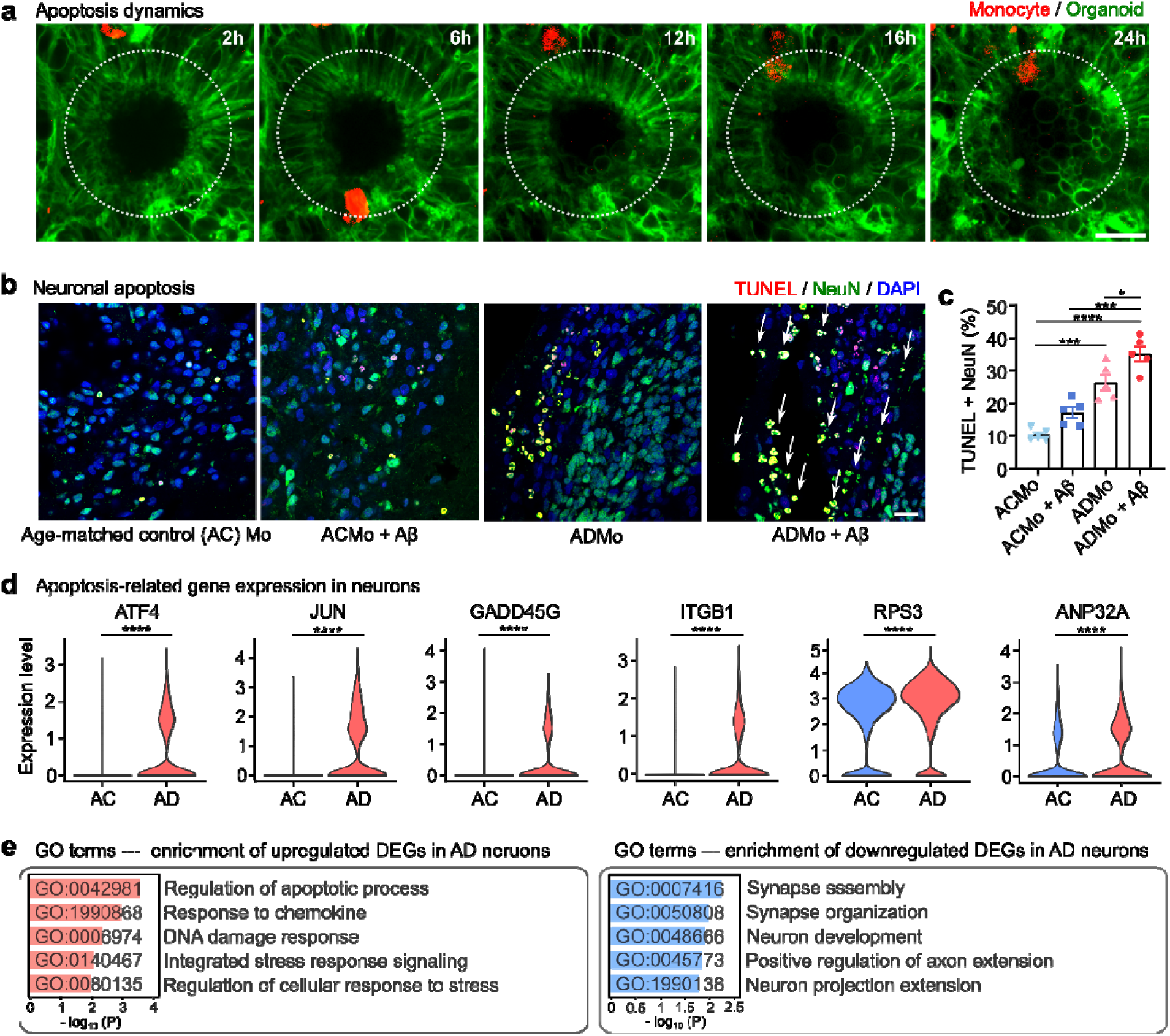
AD Monocytes (ADMo) induce neuronal apoptosis in hCO-MPSs. **(a)** Apoptosis dynamics in ADMo hCO-MPSs over 24 hours. **(b)** Representative images showing neuronal apoptosis in hCO-MPSs under different conditions: age-matched control monocytes (ACMo), ACMo + Aβ, ADMo, and ADMo + Aβ. White arrows indicate apoptotic neurons. **(c)** Quantification of neuronal apoptosis shown in (b) (mean ± s.e.m., n = 5 organoids from three independent experiments). **(d)** Expression of neuronal apoptosis-related genes in neurons within ACMo and ADMo hCO-MPSs. **(e)** Gene Ontology (GO) enrichment analysis based on neuronal DEGs in ADMo hCO-MPSs. **(a, b)** Scale bar: 20 μm.

### Increased secretion of IL-1β and CCL3 in AD monocyte-enriched hCO-MPSs

To identify the key cytokines involved in monocyte infiltration and monocyte-mediated neuroinflammation, we compared differentially expressed cytokine-encoding genes in AD hCO-MPSs and AC hCO-MPSs **(Fig. 6a**, cytokine-encoding genes are marked by triangles). We found that the pro-inflammatory cytokine IL-1β and the chemokine CCL3 were the most upregulated cytokines in AD hCO-MPSs as compared to AC hCO-MPSs. We validated our findings by ELISA assays, which consistently revealed increased protein levels of IL-1β and CCL3 in the supernatant of AD hCO-MPSs as compared to that of AC hCO-MPSs **(Fig. 6b & S8)**. As a prominent chemokine that signals via the CCR1 receptor (that we found to be highly expressed in AD monocytes) and exhibits chemotactic activity for monocytes, CCL3 may promote increased AD monocyte infiltration into the hCO-MPSs as observed in our study. Furthermore, as a central pro-inflammatory cytokine, IL-1β may contribute to monocyte-mediated neuroinflammation in the AD brain. Therefore, we propose a working model, in which CCL3 and IL-1β play the key roles in monocyte-mediated neuroinflammation in AD **(Fig. 6c)**.

**Figure 6.**
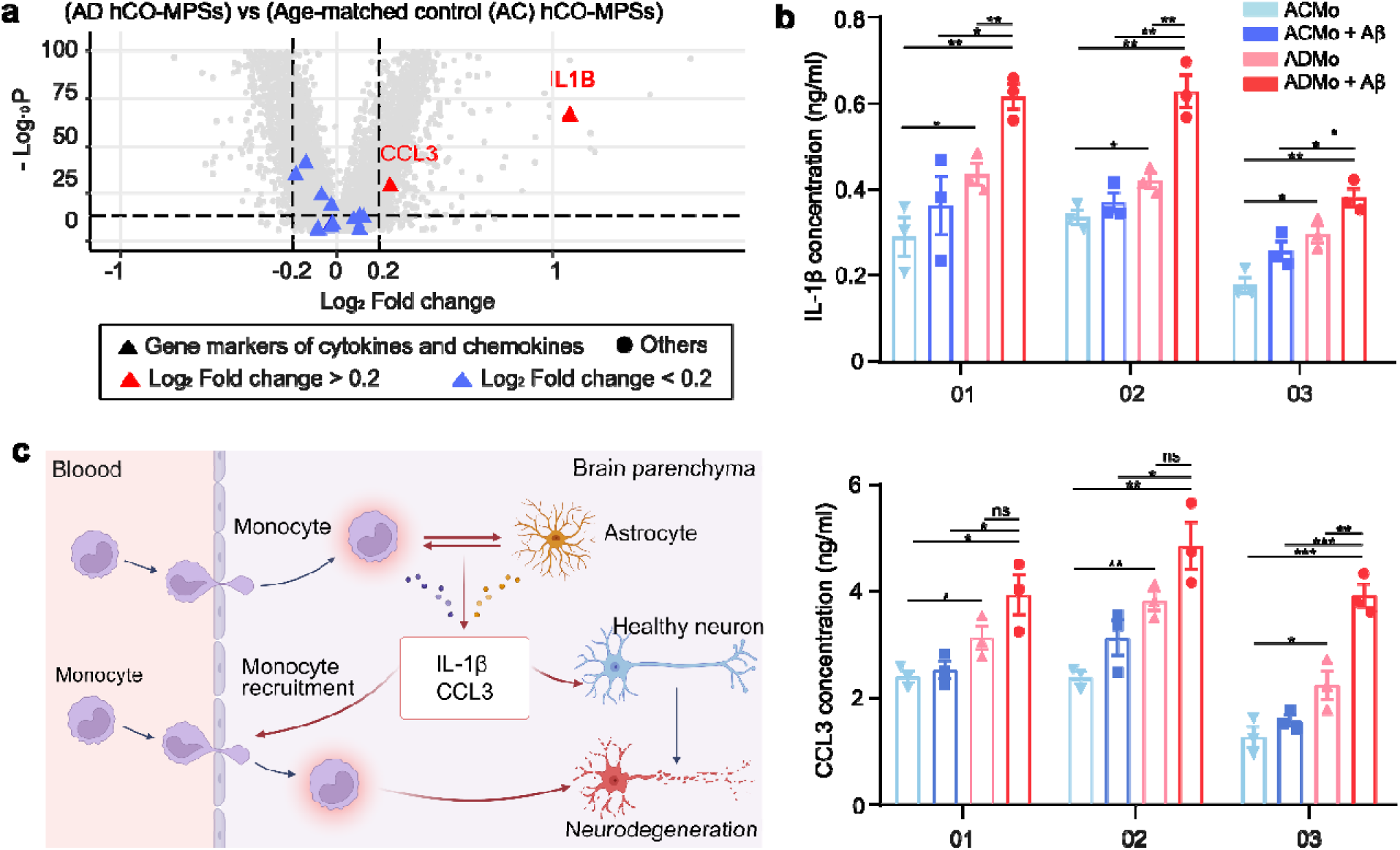
**Secreted IL-1**β **and CCL3 are increased in AD monocyte (ADMo) hCO-MPSs. (a)** A volcano plot showing differentially expressed genes (DEGs) between the ADMo and age-matched control (ACMo) hCO-MPSs. Cytokine-encoding genes were labeled with triangles and color-coded based on Log_2_ fold-change values. **(b)** ELISA assay results for IL-1β and CCL3 concentrations from 3 paired monocytes **(c)** The proposed pathway of monocyte-driven neuroinflammation.

## Discussion

In conclusion, we present the innovative hCO-MPSs as a model to explore the role of monocytes in AD neuroinflammation, a process increasingly recognized as a significant contributor to AD pathogenesis. The hCO-MPSs consisting of doughnut-shaped hCOs and 3D-printed devices were mass-produced in 96-well plates, offering advantages such as reduced necrosis and hypoxia, as well as facilitating easy infiltration by immune cells. Using this model, we investigated the dynamics of monocyte-mediated neuroinflammation in AD. Our findings revealed the pro-inflammatory alterations and effects of AD monocytes, including enhanced infiltration capacity, diminished Aβ clearance ability, stronger inflammatory response, and the induction of astrocyte activation and neuronal apoptosis, demonstrating the exacerbating role of monocytes in AD neuroinflammation. IL-1β and CCL3 were identified as key mediators in this process. Our study provides valuable insights into the impact of monocytes on AD pathogenesis, and the MPS models we developed are simple, robust, scalable, user-friendly, and compatible with current lab settings. Our MPS models have significant potential for modeling neuroinflammation, developing new therapeutics for various neuroinflammatory conditions, and contributing to the treatment of neurodegenerative diseases.

Neuroinflammation is a major driver of AD neuropathology that disrupts neuronal and glial cell homeostasis, exacerbates neurodegeneration, and likely contributes to cognitive decline^2, 4, 29, 55, 60, 64^. Using all-human cellular systems, we find that peripheral monocytes are an important source of neuroinflammation in the AD brain and induce other detrimental phenotypes, including neuronal cell death and inflammatory astrocyte activation.

Our study of peripheral monocyte infiltration is facilitated by the development of innovative doughnut-shaped hCOs. The unique hCO shape and the engineered scaffolds enhance oxygen and nutrient exchange across the organoid tissue, prevent hypoxia and necrosis of the organoid core, and support peripheral monocyte infiltration and retention within the organoid. Although other approaches, such as organoid vascularization^65^ or slicing^66^, have been applied to reduce organoid hypoxia and necrosis, our platform does not require introducing vascular cells or continued handling of individual organoids, and thus provides an easy-to-use system for differentiation of highly viable organoids. Furthermore, hCO-MPSs are compatible with time-lapse imaging, enabling the study of monocyte infiltration dynamics, whereas the 3D-printed scaffold device is easy to generate, user-friendly, and compatible with commonly used organoid differentiation protocols. We specifically chose to use cortical organoids because cortical neuropathology is evident in AD and contributes to cognitive decline^67^. Furthermore, several studies have shown that guided cortical organoid differentiation yields highly reproducible organoids as compared to unguided organoid differentiation^68–70^. Using primary human monocytes also has several advantages: (1) their human origin enables the study of human-specific disease phenotypes and molecular mechanisms; (2) monocytes derived from AD patients may contain cell-intrinsic properties specific to that patient; and (3) monocytes derived from AD patients and age-matched healthy controls may exhibit age-relevant phenotypes, given that aging is the primary risk factor for AD^71, 72^.

Our findings revealed a pro-inflammatory state of infiltrating AD monocytes, their enhanced infiltration capacity, and diminished Aβ clearance as compared to AC monocytes in hCO-MPSs. Furthermore, AD monocytes induced inflammatory astrocyte activation and neuronal apoptosis in hCO-MPSs, the phenotypes that were further exacerbated in the presence of Aβ. Taken together, these observations reveal a detrimental effect of infiltrating monocytes in AD and the potential role of monocytes in mediating neuroinflammation. We identified IL-1β and CCL3 cytokines as potential mediators of monocyte-driven neuroinflammation and recruitment of peripheral monocytes into the brain, respectively. Based on our findings, we hypothesize that, in AD, peripheral monocytes infiltrate into the brain parenchyma and interact with various brain-resident cells, especially neurons and astrocytes. These interactions contribute to an increased production of inflammatory cytokines and potent chemokines, including IL-1β and CCL3, that further promote neuroinflammation and peripheral immune cell recruitment. Notably, CCL3 is a potent chemoattractant for monocytes, and the expression of its receptor, CCR1, is also significantly upregulated in AD monocytes as compared to AC monocytes **(Fig. 2g)**. The CCL3/CCR1 axis is known to mediate monocyte recruitment during inflammation^73^, underscoring its potential role in attracting more monocytes into the brain in AD and thus further exacerbating neuroinflammation. Meanwhile, IL-1β, a pro-inflammatory cytokine and key mediator of neuronal injury^74^, may further activate astrocytes and contribute to neuronal degeneration. Thus, we propose that monocytes exacerbate neuroinflammation in AD (**Fig. 6d**). Overall, our findings provide important insights into how peripheral effectors contribute to AD progression, while our MPS platform can be applied to the development of novel therapeutics targeting neuroinflammation in future studies.

The current hCO-MPS model faces several limitations. One important thing is the absence of essential glial cells in brain organoids, such as microglia and oligodendrocytes, hindering the study of their cooperative roles in AD-related neuroinflammation. Additionally, the model lacks a functional blood-brain barrier (BBB), which plays a key role in AD neuroinflammation by regulating the passage of substances, including peripheral monocytes, into and out of the brain. Finally, other peripheral immune cells involved in AD pathology, such as T cells, B cells, and neutrophils, are also not represented in the model.

## Methods

### Human embryonic stem cell (hESC) culture

WA09 hESCs were acquired from the WiCell Research Institute and used following the regulations of WiCell and Indiana University. WA09 cells were cultured in mTeSR Plus medium (STEMCELL Technologies) on Matrigel (Corning)-coated 6-well plates. The medium was changed every two days. HESCs cells were maintained in an incubator with a constant temperature of 37°C and 5% CO_2_. Every 5 days, the cells were passaged using ReLeSR (STEMCELL Technologies).

### Generation of human cortical organoids (hCOs)

Cortical organoids were generated following the published protocol^15^. In brief, on day 0, WA09 hESCs (9,000 cells/well) were seeded in 96-well U-bottom microplates (Corning) in mTESR plus medium (STEMCELL Technologies) containing 1 μM Dorsomorphin (R&D Systems) and 10 μM SB431542 (Stemgent) to fabricate embryoid bodies (EBs). 10 µM Y-27632 (SelleckChem) was added to the induction medium for the first 24 hours. On day 3, the medium was changed to cortical organoid medium (COM) consisting of Neurobasal (Life Technologies) with 2% Gem21 NeuroPlex (Gemini Bio-Products), 1% GlutaMAX, 1% NEAA (Life Technologies), 1% N2 NeuroPlex (Gemini Bio-Products), 1% P/S (Life Technologies), 1 μM Dorsomorphin and 10 μM SB431542. COM was supplemented with 20ng/mL FGF-2 (Life Technologies) from day 10 to day 17 and with 20ng/mL FGF-2 and 20ng/mL EGF (PeproTech, Rocky Hill, NJ, USA) from day 17 to day 24. On day 24, the medium was changed to organoid maturation medium consisting of COM supplemented with 10ng/ml BDNF, 10ng/ml GDNF, 10ng/ml NT3 (all from PeproTech), 200 μM Ascorbic Acid, and 100 μM cAMP (Sigma-Aldrich) but without Dorsomorphin and SB431542. From day 30, cortical organoids were transferred to 6-well plates in COM and placed on an orbital shaker at 90 rpm. Cortical organoids were maintained with medium changes every two days.

### Design of the 3D-printed device

The device was designed using the AutoCAD software and then printed with a 3D printer (Form3B, Formlabs). The FormLabs Clear Resin V4 (FormLabs) was used as the printing material. Detailed design parameters of the device are described in **Fig. S2**.

### On-device culture of hCO-MPSs

On day 3 of cortical organoid differentiation, EBs were carefully aspirated with a 1,000 μL pipette and placed in the space between the mezzanine of the 3D-printed device and the glass bottom in a 96-well plate. 8 to 10 EBs were added into each well, and the EBs merged over time to form doughnut-shaped hCOs. After 30 days, the glass bottom was removed, and the hCO-MPSs were transferred to 6-well plate and placed on an orbital shaker at 90 rpm to enhance nutrient and oxygen exchange. The same cortical organoid differentiation protocol^15^ was used for doughnut-shaped hCOs.

### Cell viability assay

Live/dead cell staining was performed to measure the cell viability of doughnut-shaped hCOs and conventional spherical hCOs. CFSE (BioLegend) and ethidium homodimer-1 (Invitrogen) diluted at 1:1,000 were used to label live and dead cells, respectively, following the manufacturers’ protocols. Imaging was performed on an inverted fluorescence microscope (Olympus IX-83). ImageJ (version 1.54f) was used to quantify cell viability.

### Hypoxia measurement

Image-iT™ Green Hypoxia Reagent (Invitrogen) diluted at 1:1,000 was used to detect hypoxic tissue of doughnut-shaped hCOs and conventional spherical hCOs following the manufacturer’s protocol. Imaging was performed on an inverted fluorescence microscope (Olympus IX-83). ImageJ (version 1.54f) was used to quantify hypoxia fluorescence intensity.

### Whole mount staining of cortical organoids

Whole mount staining was performed to demonstrate the shape and other characteristics of doughnut-shaped hCOs. Organoids were first washed twice with 1× PBS (Gibco) and fixed with 4% paraformaldehyde at room temperature for 1 hour. Fixed organoids were washed with 1× PBS 3 times for 10 minutes, followed by incubation with organoid blocking buffer (3% fetal bovine serum, 1% (w/v) bovine serum albumin, 0.5% Triton X-100, 0.5% Tween 20, and 0.01% (wt/vol) sodium deoxycholate in 1× PBS) at room temperature for 2 hours. After blocking, the organoids were incubated with primary antibodies (diluted in the blocking buffer) with gentle rocking at 4°C overnight. The organoids were then washed with PBST (0.1% Tween 20 in 1× PBS) 5 times for 5 minutes and incubated with secondary antibodies and DAPI (Invitrogen) (diluted in the blocking buffer) with gentle rocking and protected from light at 4°C overnight. After incubation with secondary antibodies, the organoids were washed with PBST 5 times for 5 minutes and once with 1× PBS followed by dehydration with 50%, 70%, and 100% methanol or ethanol for 10 minutes each. Murray’s clear (2:1 benzyl benzoate-to-benzyl alcohol) was used for tissue clearing of the dehydrated organoids. Detailed information of antibodies and their dilutions used in this study is provided in Table S1.

### Cryo-sectioning of organoids

For cryo-sectioning of hCOs, organoids were washed twice with 1× PBS and fixed with 4% paraformaldehyde at 4°C overnight followed by dehydration in 30% sucrose in 1× PBS (w/v) at 4°C overnight. Organoids were then transferred to a cryomold (Sakura Finetek) with O.C.T. compound (Fisher Scientific) and frozen on dry ice. Embedded organoids were sectioned using a cryostat (Leica), and 30 μm-thick slices were collected on Superfrost Plus slides (VWR International).

### Immunofluorescence staining

The slides of cryosectioned organoids were washed with 1× PBS twice for 5 minutes to remove the O.C.T compound and submerged in the Antigen Retrieval Solution (Thermo Fisher) and boiled for 20 minutes. After the slides cooled down, they were rinsed with 1× PBS and incubated with a blocking buffer (5% fetal bovine serum and 0.3% Triton™ X-100 in 1× PBS) for 1 hour. Subsequently, the slides were incubated with primary antibodies (diluted in the blocking buffer) protected from light at 4°C overnight. After incubation, the slides were washed with 1× PBS 5 times for 10 minutes, followed by incubation with secondary antibodies (diluted in the blocking buffer) protected from light at room temperature for 2 hours. Finally, the slides were washed with 1× PBS 3 times for 10 minutes and incubated with DAPI diluted in 1× PBS for 10 minutes. Detailed information of antibodies used in the experiment can be found in Table S1.

### THP-1 cell culture

The THP-1 cells (ATCC TIB-202) were used in preliminary experiments to establish the co-culture system before using patients’ monocytes. The culture medium for THP-1 cells was RPMI 1640 (Gibco) supplemented with 10% fetal bovine serum (Gibco), 0.05 mM 2-mercaptoethanol (Sigma), and 100 U/mL penicillin/streptomycin (Gibco). The cells were maintained in a humidity incubator at 37°C temperature and 5% CO_2_. The cells were passaged when the cell density reached 1 × 10^6^ cells/mL. The culture medium was changed every two days.

### Time-lapse imaging

The dynamic processes of monocyte infiltration and cell apoptosis were recorded using time-lapse microscopy. Cortical organoids expressing ChR2-EYFP (derived from H9-CAG-ChR2-EYFP hESCs) were co-cultured with monocytes (1×10^5^ monocytes/organoid) labeled with the Vybrant Dil solution (Invitrogen) for time-lapse imaging experiments. Time-lapse images were taken at regular intervals using the Olympus OSR Spinning Disk confocal microscope. Videos were created with Adobe Photoshop 2024.

### Monocyte isolation from human peripheral blood mononuclear cells (PBMCs)

PBMCs of AD patients were isolated from whole blood obtained from Indiana University Medical School using SepMate™-50 (IVD) (STEMCELL Technologies) following the manufacturer’s protocol. PBMCs of age-matched control donors were purchased from STEMCELL Technologies. Monocytes were isolated by positive selection using CD14 MicroBeads UltraPure and MS Columns (Miltenyi Biotec Inc) following the manufacturer’s protocol. Typically, 1.5×10^5^ monocytes can be isolated from 1×10^6^ PBMCs. Demographic information of AD patients and age-matched control donors is provided in Table S2.

### Imagining monocyte infiltration

To study monocyte infiltration, monocytes were labeled with the Vybrant Dil solution (Invitrogen) in serum-free RPMI-1640 (Gibco) and incubated at 37°C and 5% CO_2_ for 30 minutes. Labeled monocytes were washed twice, and 1×10^5^ monocytes were seeded onto each hCO-MPS. In Aβ treatment experiments, hCO-MPSs were pre-incubated with 1 μM of HyLite 488 labeled Aβ (1-42) (AnaSpec) and incubated on a shaker overnight to form Aβ aggregates. Monocyte infiltration was imaged using the Olympus OSR Spinning Disk confocal microscope inside a Tokai Hit on-stage incubator set at 37°C and 5% CO_2_. Quantification was performed using Imaris (9.0.1).

### Preparation of Aβ aggregates

Beta-Amyloid (1-42) peptides (AnaSpec) were resuspended in dimethylsulfoxide (DMSO) to 10 mM and sonicated for 10 minutes to form Aβ aggregates. Aβ aggregates were diluted to 1 mM in 1× PBS and stored at -80°C. Immediately prior to use, Aβ aggregates were diluted to the final concentration used in experiments.

### Aβ phagocytosis

The hCO-MPSs were loaded with HyLite 488 Aβ (1-42) (AnaSpec) with overnight rocking. The DiL membrane tracker (Invitrogen) labeled monocytes were co-cultured with Aβ treated organoids for 24 hours before visualization using a Leica SP8 microscope. Quantification of Aβ phagocytosis was performed using ImageJ (v 1.54f).

### Sample preparation for single-cell RNA sequencing

48 hours after monocyte co-culture with hCO-MPSs, AD and AC hCO-MPSs were dissociated into a single-cell suspension using a papain-based dissociation kit (Neural Tissue Dissociation Kit-P, Miltenyi Biotec), with slight modifications to the manufacturer’s protocol. Specifically, organoids were washed three times with HBSS without Ca²⁺ and Mg²⁺ and cut into hemispheres using scissors. The hemispheres were incubated in enzyme mix 1 (enzyme P) with orbital shaking at 90 rpm for 15 minutes. Subsequently, enzyme mix 2 (enzyme A) was added, and the incubation continued for an additional 30 minutes with periodic trituration every 10 to 15 minutes using a wide-bore 1 mL pipette tip. After incubation, the samples were filtered through a 30 μm strainer (Miltenyi Biotec) to remove large aggregates and retain single cells. The resulting single-cell suspension was centrifuged at 300 x g for 8 minutes at 4°C, resuspended in organoid medium without growth factors to a concentration of 1×10^6^ cells/mL, and subjected to scRNA-seq (10x Genomics).

### ScRNA-seq data analysis

Raw sequencing data were preprocessed using the Cell Ranger software on the 10x Genomics cloud platform (v6.1.2). Reads were aligned and quantified using the hg38 human reference genome (refdata-gex-GRCh38-2020-A, 10x Genomics). Downstream data analysis was performed in R (v4.3.2) using the Seurat package (v4.4.0)^75^. Genes expressed in less than 10 cells were excluded to remove low quality genes; cells with nFeature_RNA between 200 to 7,000, nCount_RNA less than 25,000, and percent.mt less than 10% were kept in the dataset for downstream analysis. The data were analyzed following a standard workflow^76^. Briefly, normalization and feature selection were performed for the Seurat object, and the FindIntegrationAnchors and IntegrateData Seurat functions were used to integrate the datasets. The output was then passed through ScaleData, RunPCA, FindNeighbors, and FindClusters functions in Seurat for data scaling, linear dimensional reduction, and cell clustering. Uniform Manifold Approximation and Projection (UMAP) was applied for data visualization. The cells projected on the UMAP space were annotated using canonical marker genes for neurons, progenitors, intermediate progenitors, astrocytes, other glial cells, and monocytes^15, 77, 78^. Differentially expressed genes (DEGs) between cell clusters and groups were identified using the FindMarkers function in Seurat. Only DEGs with a p value of < 0.05 were included in the analysis.

### Gene ontology (GO) enrichment analysis

DEGs with a p value of < 0.05 were imported into the Enricher analysis platform (https://maayanlab.cloud/Enrichr/) for GO analysis according to the instructions^79^. Only GO terms with a p value of < 0.05 were included in the analysis.

### TUNEL staining

Terminal deoxynucleotidyl transferase dUTP nick end labeling (TUNEL) staining was performed on cryosectioned organoids to visualize cell apoptosis using the In Situ Cell Death Detection Kit (Roche) according to the manufacturer’s protocol. Briefly, the slides were first fixed in fixation solution (4% Paraformaldehyde in 1× PBS) at room temperature for 20 minutes followed by washing with 1× PBS for 30 minutes. Subsequently, the slides were incubated with the permeabilization solution (0.1% Triton X–100 and 0.1% sodium citrate in 1× PBS) on ice for 2 minutes. The slides were then incubated with the TUNEL reaction mixture at 37°C in a dark humidified box for 1 hour. Finally, the slides were mounted with the antifade mounting medium. Imaging was performed using a confocal microscope (Leica Stellaris 8). Quantification of neuronal apoptosis was performed in ImageJ (v 1.54f).

### ELISA assay

The enzyme-linked immunosorbent assay (ELISA) was used to measure cytokine concentrations in the cell culture supernatant. 48 hours after monocyte co-culture with hCO-MPSs, 250 μL of supernatant was aspirated with a pipette and centrifuged at 300 g for 10 minutes to remove cell debris. 200 μL of the supernatant was then diluted 3 and 10 times for the measurement of IL-1β and CCL3 concentrations, respectively. Human IL-1 beta ELISA Kit and Human MIP-1α (CCL3) ELISA Kit (Invitrogen) were used following the manufacturer’s protocol. For each ELISA assay test, 50 μL of the diluted supernatant was used, and the absorption reading was performed on the Synergy H1 plate reader (BioTek).

### RT-qPCR analysis

To analyze gene expression profiles of hCO-MPSs, the organoids were first washed twice with 1× PBS, and the RNA was extracted using the Quick RNA MicroPrep Kit (Zymo). The extracted RNA was immediately reverse transcribed into complementary DNA (cDNA) using the qScript cDNA synthesis kit (Quantabio). The resulting cDNA was then subjected to qPCR analysis using the SYBR Green real-time PCR master mix (Thermo Fisher). Detailed primer sequences used for qPCR are listed in Table S3. Data were analyzed using the ΔΔCt method, where the delta Ct value was calculated by subtracting the Ct value of the target gene in day 1 embryoid bodies (EB) from that in the organoids. The ΔCt values were normalized to the housekeeping gene GAPDH. Each qPCR reaction was performed in triplicate.

### ROS measurement

To detect ROS production, monocytes were labeled with the Vybrant DiO dye (Invitrogen) diluted 1:1,000 in RPMI-1640 for 1 hour. Monocytes were then loaded onto the organoid device or cultured with Aβ (1-42) (AnaSpec). The ROS production was then detected with the CellROX reagent (Gibco) that was added into the co-culture system at 5 µM for 30 min. The cells were then immediately fixed with 10% formalin and washed twice with 1× DPBS (Gibco). The ROS dye was visualized under an Olympus IX-83 inverted fluorescent microscope, and the data analysis was performed using ImageJ (v 1.54f).

### Statistical analysis

The Kolmogorov-Smirnov test was used to assess normality, while the F-test was used to compare variances. Unpaired t-tests were used for the statistical comparison of experimental groups. The statistical significance of differences in values is denoted as: *p<0.05, **p<0.01, ***p<0.005. ****p<0.001. All statistical analyses were performed in GraphPad Prism 8.

## Acknowledgments

We acknowledge Drs. Z. Li, and N. Wang for their discussions and help on data analysis and sample preparation. The work was partially supported by the National Institute of Health Awards (DP2AI160242, and U01DA056242). The authors also thank the Indiana University Imaging Center (NIH1S10OD024988-01) for providing their instruments. J.C. is a predoctoral scholar in the Stem Cell Biology and Regenerative Medicine Research Training Program of the California Institute for Regenerative Medicine (CIRM).

